# Deep mutational scanning of SARS-CoV-2 Omicron BA.2.86 and epistatic emergence of the KP.3 variant

**DOI:** 10.1101/2024.07.23.604853

**Authors:** Ashley L. Taylor, Tyler N. Starr

## Abstract

Deep mutational scanning experiments aid in the surveillance and forecasting of viral evolution by providing prospective measurements of mutational effects on viral traits, but epistatic shifts in the impacts of mutations can hinder viral forecasting when measurements were made in outdated strain backgrounds. Here, we report measurements of the impact of all single amino acid mutations on ACE2-binding affinity and protein folding and expression in the SARS-CoV-2 Omicron BA.2.86 spike receptor-binding domain (RBD). As with other SARS-CoV-2 variants, we find a plastic and evolvable basis for receptor binding, with many mutations at the ACE2 interface maintaining or even improving ACE2-binding affinity. Despite its large genetic divergence, mutational effects in BA.2.86 have not diverged greatly from those measured in its Omicron BA.2 ancestor. However, we do identify strong positive epistasis among subsequent mutations that have accrued in BA.2.86 descendants. Specifically, the Q493E mutation that decreased ACE2-binding affinity in all previous SARS-CoV-2 backgrounds is reversed in sign to enhance human ACE2-binding affinity when coupled with L455S and F456L in the currently emerging KP.3 variant. Our results point to a modest degree of epistatic drift in mutational effects during recent SARS-CoV-2 evolution but highlight how these small epistatic shifts can have important consequences for the emergence of new SARS-CoV-2 variants.

## INTRODUCTION

The evolution of SARS-CoV-2 is marked by the continuous emergence of viral variants (Carabelli et al. 2023). Mutations in these variants often concentrate within the spike protein and in particular its receptor-binding domain (RBD), where they enable escape from neutralizing antibody immunity while maintaining or improving ACE2 receptor binding and cellular entry (Maher et al. 2022; Ma et al. 2023). Though advances in genomic sequencing capacity have enabled real-time surveillance of the origin and spread of these viral variants, it remains difficult to rapidly determine the functional consequences of the mutations they sample.

Deep mutational scanning (Fowler & Fields 2014) has emerged as a powerful experimental method to aid in viral surveillance. These experiments comprehensively measure the impacts of all single amino acid changes in the SARS-CoV-2 spike or RBD on key phenotypes such as receptor-binding affinity, cellular entry, and antibody escape (Cao et al. 2022; Dadonaite et al. 2024; Francino-Urdaniz et al. 2021; Greaney et al. 2021; Kugathasan et al. 2023; Moulana et al. 2022; Ouyang et al. 2022; Starr et al. 2020; Starr, Greaney, Addetia, et al. 2021; Starr, Greaney, Dingens, et al. 2021; Taft et al. 2022). Because these measurements are made prospectively (i.e. prior to the evolution of a variant), they can be immediately consulted upon discovery of a novel viral variant to provide preliminary insights into the properties it might exhibit. Deep mutational scanning can therefore provide early insights about a novel variant while more intensive retrospective analyses of variant phenotype proceed. More recently, analysis of large deep mutational scanning datasets also shows promise in the forecasting and prediction of future viral evolution (Cao et al. 2023; Dadonaite et al. 2024; Greaney, Starr, & Bloom 2022; Maher et al. 2022).

However, the impacts of mutations on protein function are not constant over time due to epistasis, the phenomenon where the functional impact of one mutation is modulated by another (Starr & Thornton 2016). We and others have described important epistatic interactions shaping SARS-CoV-2 evolution, such as the interaction between N501Y and Q498R that facilitated the original emergence of Omicron (Moulana et al. 2022; Starr, Greaney, Hannon, et al. 2022; Zahradník et al. 2021) and interactions between substitutions at positions 455, 456, and/or 493 that have shaped subsequent Omicron evolution (Jian et al. 2023, 2024; Taylor & Starr 2023). The utility of deep mutational scanning data in SARS-CoV-2 surveillance and forecasting therefore depends on updated measurements of mutational effects within each major variant that emerges, as the effects of mutations measured in an earlier variant may not accurately portray the impact that mutation would have in the viruses that circulate at present.

Since the original emergence of Omicron BA.1 in November, 2021, a string of derivative Omicron lineages (e.g., BA.2, BA.4/BA.5, XBB.1.5) have continued to evolve and displace prior strains in a more or less linear fashion. However, in August, 2023, a new “saltation” variant dubbed BA.2.86 gathered attention due to its large number of sequence substitutions relative to concurrently circulating strains and its rapid detection across multiple countries around the world suggesting early spread. BA.2.86 and the predominant strains in August, 2023 such as XBB.1.5, EG.5 (XBB.1.5 + F456L) or HK.3 (EG.5 + L455F, i.e. “FLip”) last shared a common ancestor in BA.2, meaning BA.2.86 had been evolving for >1 year separately from the competing strains, allowing it to sample unique mutations such as a single amino acid deletion at position 483 (near major antibody epitopes) and the presence of a novel glycan at position 354.

Due to the large number of sequence substitutions in BA.2.86 (34 spike mutations relative to BA.2, and 14 in the RBD, **Figure 1A**), there was immediate concern whether it would have a similar advantage in antibody-escape over competing XBB.1.5 variants as the original Omicron strain exhibited compared to pre-Omicron variants. However, BA.2.86 was soon found to exhibit a similar degree of resistance to serum antibodies as the competing XBB.1.5 descendants (Khan et al. 2023; Liu et al. 2024; Wang et al. 2023; Yang et al. 2023). Nonetheless, BA.2.86 did show a slight transmission advantage over co-circulating strains (Tamura et al. 2024), and following acquisition of the L455S mutation in JN.1 that further optimized antibody evasion (Yang et al. 2024), rose to dominate case counts in December, 2023. JN.1 is now spinning out further derivatives (e.g., KP.3) exhibiting enhanced fitness (Jian et al. 2024; Kaku, Uriu, et al. 2024; Kaku, Yo, et al. 2024), suggesting future SARS-CoV-2 evolution will continue for some time to stem from the BA.2.86 evolutionary lineage.

**Figure 1.**
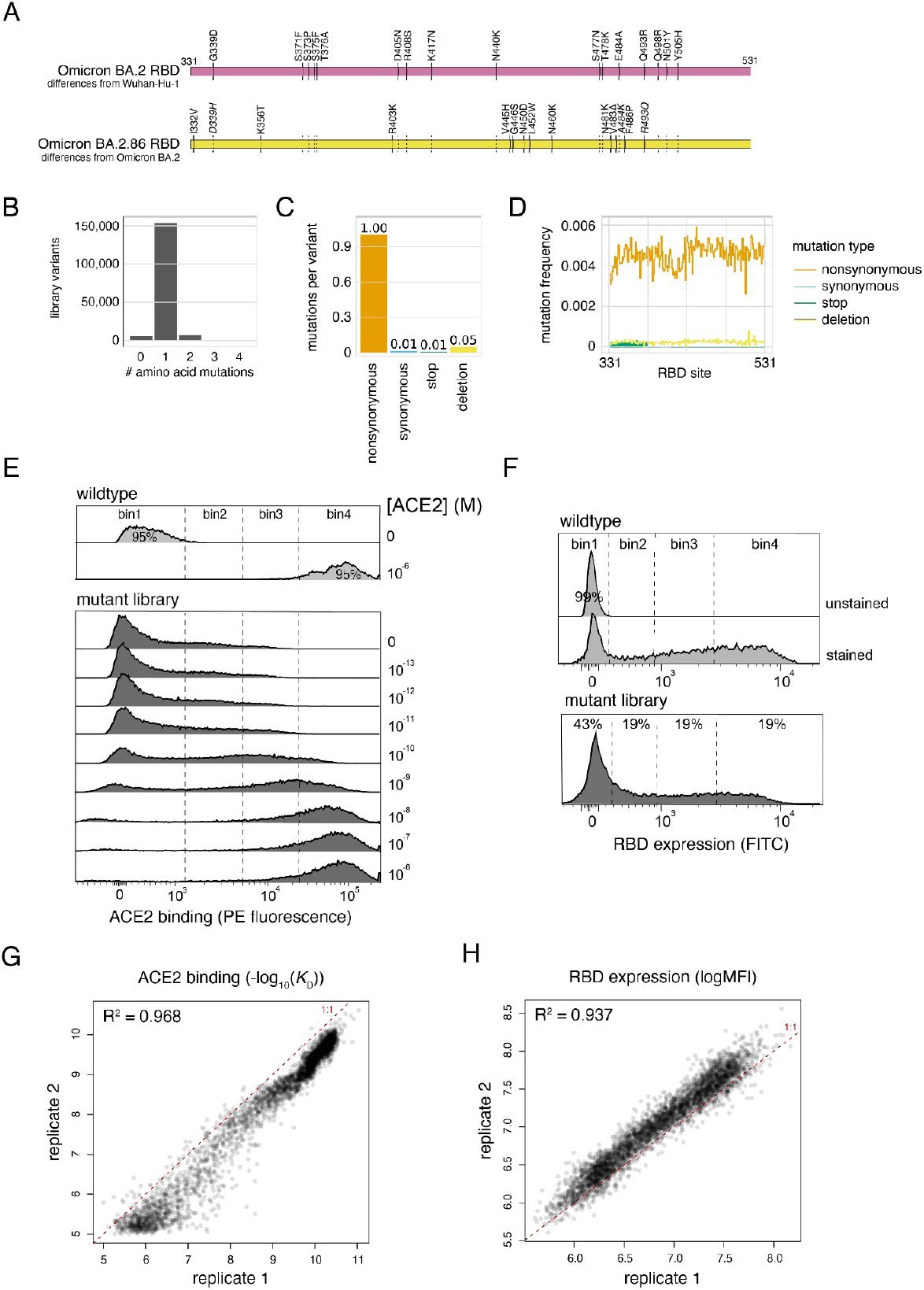
Deep mutational scanning of the SARS-CoV-2 Omicron BA.2.86 RBD. (**A**) Diagram of the RBD substitutions that distinguish Omicron BA.2 from Wuhan-Hu-1 (top), and BA.2.86 from BA.2 (bottom). Italicized mutations in BA.2.86 indicate secondarily mutated (D339H, A484K) or reverted (R493Q) substitutions that originally changed from Wuhan-Hu-1, and dashed lines show propagation of BA.2. changes to BA.2.86. Wuhan-Hu-1 reference spike numbering is used throughout the manuscript. (**B-D**) Quality control of the BA.2.86 RBD site-saturation mutagenesis library as assessed by PacBio sequencing, illustrating the distribution of number of amino acid mutations per barcoded variant (B), the average number of mutations of each type across library variants (C), and the distribution of mutations across sites in the RBD over all variants (D). (**E, F**) FACS gates used to sort RBD^+^ singlet cells for ACE2 titration (E) and RBD expression (F) deep mutational scanning experiments from one representative replicate. (**G, H**) Correlation in per-mutant deep mutational scanning measurements between independently barcoded replicate libraries for ACE2-binding affinity (G) and RBD expression (H) experiments.

To aid in ongoing viral surveillance and forecasting, here we report deep mutational scanning data in the BA.2.86 RBD. We measure the impacts of all single amino acid changes and single-codon deletions in the BA.2.86 RBD on ACE2-binding affinity and RBD folding/expression, revealing continued tolerance to mutation within this viral domain. We find that mutational effects in BA.2.86 largely resemble those as measured in BA.2 despite their large sequence divergence. Nonetheless, we find a key epistatic interaction between Q493E and mutations at positions 455 and 456 that support the currently emerging KP.3 sub-variant.

## RESULTS

### Deep mutational scanning of the Omicron BA.2.86 RBD

We have previously described a yeast-surface display platform for deep mutational scanning of the SARS-CoV-2 RBD (Adams et al. 2016; Starr et al. 2020). We have used this platform to measure the impacts of mutations in the Wuhan-Hu-1, Alpha, Beta, Delta, Eta, BA.1, BA.2, BQ.1.1, and XBB.1.5 strains on ACE2-binding affinity and RBD folding efficiency (Starr, Greaney, Hannon, et al. 2022; Starr, Greaney, Stewart, et al. 2022; Taylor & Starr 2023). Here, we follow this pipeline to generate mutational data in the SARS-CoV-2 BA.2.86 variant RBD background.

We first created a site-saturation mutagenesis library introducing every possible single amino acid mutation across the BA.2.86 RBD. We also programmed single-codon deletions at each position, as well as premature stop codons or silent encoding of the wildtype codon at a fraction of sites as internal controls. We cloned library mutants together with an variant-identifier N16 nucleotide barcode, and we used PacBio long-read sequencing to link library barcodes with their associated RBD mutant and survey the composition of our library. Our library showed the expected balance of single-mutant variants (**Figure 1B**) of the expected types (**Figure 1C**), with even mutation rates across the 200 RBD positions (**Figure 1D**).

We transformed the BA.2.86 RBD mutant library into a yeast-surface display platform that enables genotype-phenotype linkage between the mutant plasmid a yeast contains and the folded RBD protein that it expresses tethered to its cell surface. We incubated yeast-displayed RBD libraries across a concentration gradient of fluorescently-labeled monomeric human ACE2 or an antibody targeting a C-terminal c-Myc tag, and we used fluorescence-activated cell sorting (FACS) to partition library variants on the basis of activity (**Figure 1E,F**). We deep sequenced the linked barcodes in each FACS bin and determined the impact of each mutation on ACE2-binding affinity or RBD expression level based on the distribution of sequencing counts across FACS bins. These deep mutational scanning measurements were conducted in duplicate with independently barcoded variant libraries, with per-mutant phenotypes strongly correlated between replicates (**Figures 1G, H**).

Heatmaps illustrating the impact of each RBD mutation on ACE2-binding affinity and RBD expression are presented in **Figure 2**. To aid visualization of these large datasets, interactive heatmaps, including measurements from prior SARS-CoV-2 variants, are available at https://tstarrlab.github.io/SARS-CoV-2-RBD_DMS_Omicron-EG5-FLip-BA286/RBD-heatmaps/.

**Figure 2.**
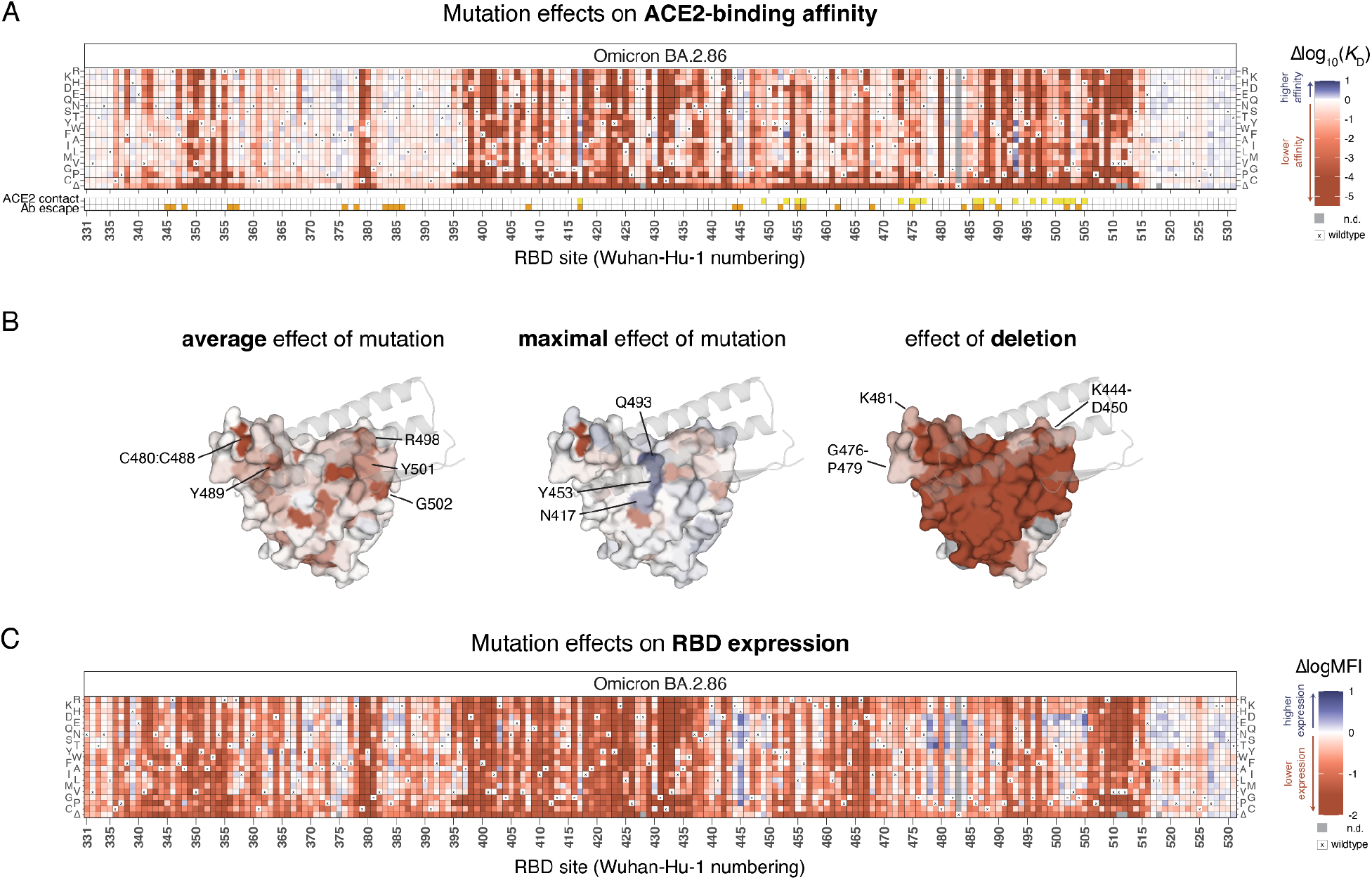
Effects of mutations in the BA.2.86 receptor-binding domain on ACE2-binding and RBD expression. (**A**) Heatmap illustrating the impacts of all mutations in the BA.2.86 RBD on ACE2-binding affinity as determined from FACS-seq experiments with yeast-displayed RBD mutant libraries. ACE2 contact residues (yellow squares, bottom) defined as RBD residues with non-hydrogen atoms <5Å from ACE2 in the BA.2.86 RBD structure (PDB 8QSQ; Liu et al. 2024). Antibody escape residues (orange squares, bottom) defined as those with average >0.125 relative antibody escape from aggregated deep mutational scanning data (Greaney, Starr, & Bloom 2022). (**B**) Deep mutational scanning data from (A) mapped to the ACE2-bound BA.2.86 RBD structure (PDB 8QSQ; Liu et al. 2024), illustrating the average effect of mutations at a site (left), the maximal effect of any mutation at a site (center), or the effect of the single-codon deletion (right). Sites of interest are labeled, and ACE2 (key motifs only) is shown as transparent gray cartoon. (**C**) Heatmap illustrating the impacts of all mutations in the BA.2.86 RBD on yeast-surface expression levels, a proxy for folding and expression efficiency. Individual measurements from (A) and (C) are reported in Supplemental Data S1, and an interactive version of these heatmaps is available at https://tstarrlab.github.io/SARS-CoV-2-RBD_DMS_Omicron-EG5-FLip-BA286/RBD-heatmaps/.

As with deep mutational scans of prior SARS-CoV-2 variants, we find considerable tolerance to mutation in the RBD, with many amino acid mutations having neutral or positive impacts on ACE2-binding affinity, including those that directly contact ACE2 (indicated by yellow squares, **Figure 2A**). Although some direct contact residues are highly constrained (e.g., Y489 [Wuhan-Hu-1 reference numbering used throughout], R498, Y501, and G502; **Figure 2B**, left), many contact residues are tolerant to mutation and can sample affinity-enhancing mutations (e.g., N417, Y453, Q493; **Figure 2B**, center). Deletions at the ACE2 interface are primarily deleterious, though two peripheral loops where indels occur more broadly during sarbecovirus evolution are less constrained (**Figure 2B**, right) (Taylor & Starr 2023).

Our experiments also identify mutations that would enhance RBD expression (**Figure 2C**), a proxy for folding and stability (Kowalski et al. 1998; Shusta et al. 1999). Many of these stabilizing mutations remove basic residues at the ACE2 interface that have been introduced in variant evolution since Wuhan-Hu-1, for example mutations to H445 (originally V445 in Wuhan-Hu-1), K478 (originally T478), K481 (originally N481), and K484 (originally E484). The increase in positively charged residues at the ACE2 interface conferred by these basic BA.2.86 residues has been noted to increase electrostatic complementarity with the corresponding ACE2 surface that is rich in acidic residues (Wang et al. 2023), which our data suggest may come at some cost for isolated RBD folding efficiency. We previously identified stabilizing space-filling mutations (Starr et al. 2020) that compensate for a loss of lipid tail volume that occupies space in the RBD core in the locked spike trimer structure (Toelzer et al. 2020); our deep mutational scanning map in BA.2.86 shows similar patterns of space-filling stabilizing mutations in the RBD core (e.g., I358F, A363Y), suggesting that the positive impact that these mutations had on RBD vaccine stability in earlier variants (Ellis et al. 2021) will continue in vaccine designs derived from BA.2.86.

### Epistatic shifts in mutational effects

We next determined the extent to which the effects of mutations in the BA.2.86 RBD differ compared to their effects in the BA.2 ancestor due to epistasis. We computed a sitewise “epistatic shift” metric for each RBD residue (Starr, Greaney, Hannon, et al. 2022), which identifies sites where the effects of mutations on ACE2-binding affinity differ more or less strongly between a pair of variants (**Figure 3**; interactive visualization of all variants and sites available at https://tstarrlab.github.io/SARS-CoV-2-RBD_DMS_Omicron-EG5-FLip-BA286/epistatic-shifts/). The epistatic shift is a probabilistic distance metric (Jensen-Shannon distance) that compares sitewise profiles of mutational effects between two variant backgrounds, computed on the vectors of affinities measured for the 20 amino acids possible at a position. The epistatic shift metrics scales from 0 to 1, with 0 indicating a site where the measured affinities of each amino acid mutant are identical between backgrounds and 1 indicating a site where the distributions are entirely dissimilar.

**Figure 3.**
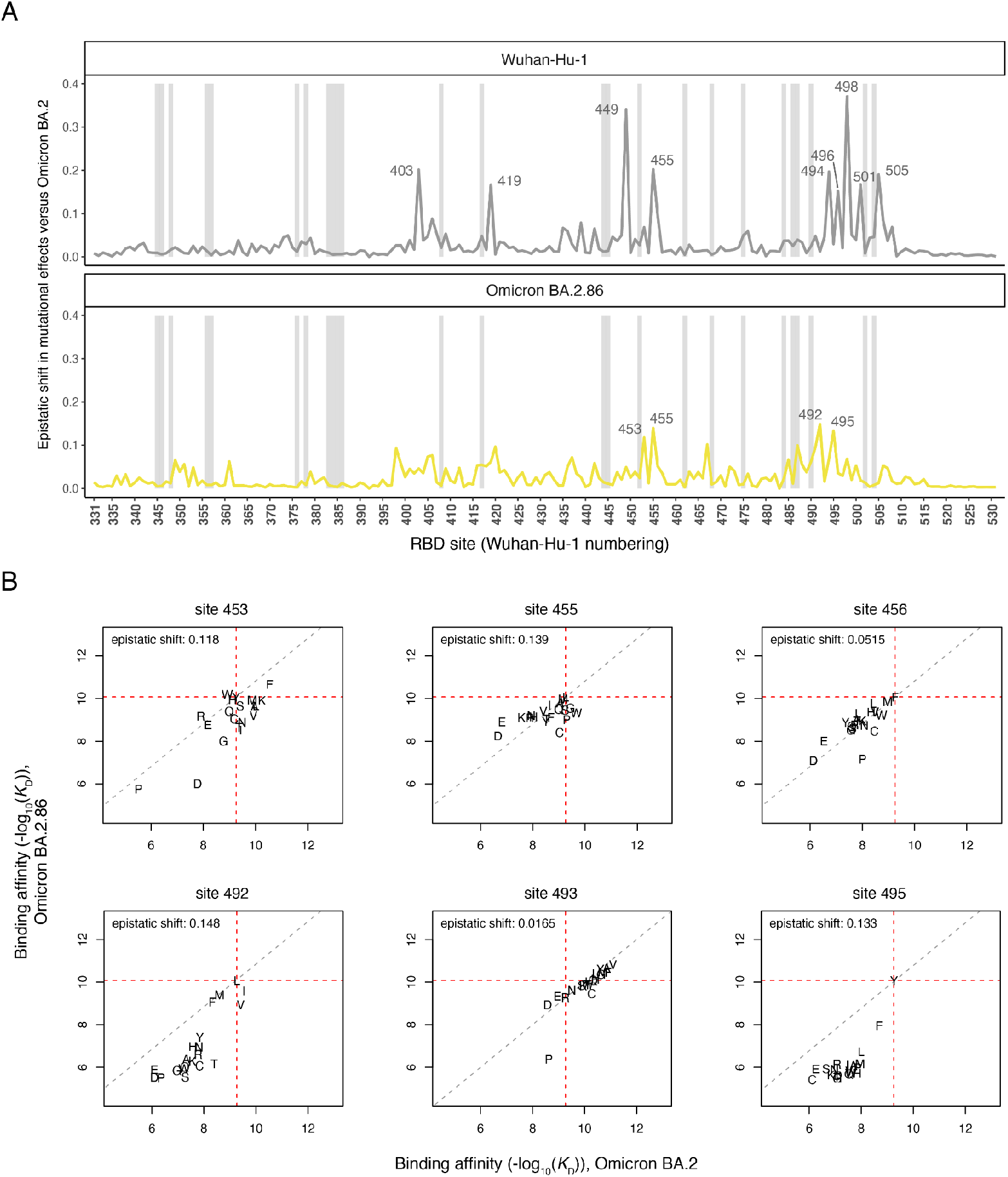
Epistatic shifts in mutational effects on ACE2 binding. (**A**) Epistatic shift in the effects of mutations on ACE2 binding at each RBD position as measured in the Wuhan-Hu-1 (previously reported in (Starr, Greaney, Hannon, et al. 2022)) or BA.2.86 background compared to those previously measured in Omicron BA.2 (Starr, Greaney, Stewart, et al. 2022). Gray bars indicate sites of strong antibody escape, as defined in Fig. 2A. (**B**) Mutation-level plots of epistatic shifts between BA.2 and BA.2.86 at sites of interest. Each scatterplot shows the measured ACE2-binding affinity of each amino acid (plotting character) in the BA.2.86 versus BA.2. backgrounds. Red dashed lines mark the wildtype RBD affinities on each axis, and the gray dashed line indicates the additive (non-epistatic) expectation. Interactive plots enabling the comparison of all SARS-CoV-2 variants and scatterplots for all RBD sites are available at https://tstarrlab.github.io/SARS-CoV-2-RBD_DMS_Omicron-EG5-FLip-BA286/epistatic-shifts/.

We previously found that the 16 substitutions separating Omicron BA.2 from the ancestral Wuhan-Hu-1 variant induced strong and widespread epistatic shifts across the RBD’s ACE2-contact interface (**Figure 3A**, top), largely attributed to the impact of the N501Y substitution (Starr, Greaney, Hannon, et al. 2022; Starr, Greaney, Stewart, et al. 2022). BA.2.86 is separated from BA.2 by a similar number of RBD changes (14 changes) as BA.2 itself is separated from Wuhan-Hu-1 (**Figure 1A**), raising the question whether similarly dramatic epistatic shifts are present between BA.2 and BA.2.86. The sitewise epistatic shifts between BA.2.86 and BA.2, however, are notably modest compared to the epistatic shifts seen between Wuhan-Hu-1 and BA.2 (**Figure 3A**, bottom). Among the sites with the strongest epistatic shifts (**Figure 3B**), the more common trend is that mutations that enhanced ACE2 affinity in the BA.2 background become constrained in BA.2.86 (lower-right quadrant of scatterplots in **Figure 3B**, e.g. V453, W455, or I492) instead of mutations that were deleterious in BA.2 becoming affinity-enhancing in BA.2.86 (upper-right quadrant, e.g. W453). Despite showing few newly expanded pathways for ACE2-affinity enhancement, BA.2.86 itself has been found to have tighter binding affinity for ACE2 compared to BA.2 and competing XBB.1.5 subvariants (Liu et al. 2024; Wang et al. 2023; Yang et al. 2023), which also expands pathways of evolution by offsetting mutations like L455S that trade off improved antibody escape with slightly reduced ACE2-binding affinity (Jian et al. 2024; Yang et al. 2024).

### Epistatic emergence of the KP.3 variant

Following the initial discovery of BA.2.86, its rise in global frequency was relatively slow until derivative lineages acquired additional spike changes that increased fitness. The first BA.2.86 descendant to rise to global dominance and displace the competing XBB.1.5-related variants was JN.1 (BA.2.86 + L455S) (Jian et al. 2024; Yang et al. 2024), and as of July, 2024, KP.3 variants which also sample F456L and Q493E (**Figure 4A**) comprise the majority of global sequences (Kaku, Uriu, et al. 2024; Kaku, Yo, et al. 2024). The Q493E mutation in KP.3 presented as a particular surprise, as this mutation has been strongly deleterious to ACE2 binding throughout SARS-CoV-2 variant evolution, decreasing affinity by approximately one order of magnitude in previously measured backgrounds and decreasing affinity ∼5.4-fold in the BA.2.86 data reported here.

**Figure 4.**
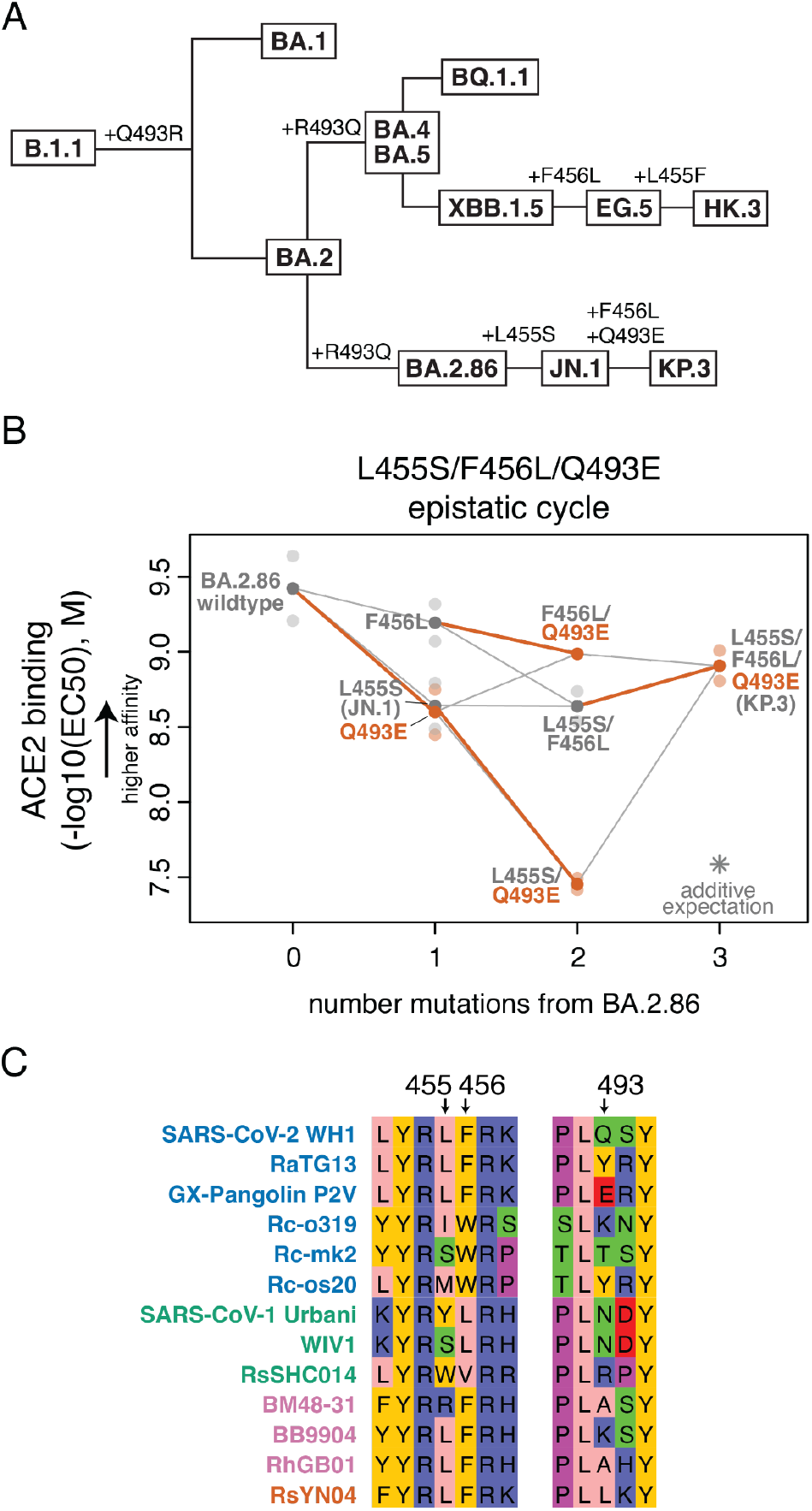
Epistatic emergence of the KP.3 variant. (**A**) Cladogram showing relationships among select SARS-CoV-2 Omicron variants, with amino acid substitutions at positions 455, 456, and 493 indicated (other mutations not shown). (**B**) Triple mutant cycle diagram illustrating epistatic interactions between L455S, F456L, and Q493E underlying KP.3 variant evolution. Transparent points indicate duplicate measurements of each variant’s binding strength for human ACE2 (determined as the EC50 from titrations of monomeric human ACE2 over yeast-displayed RBD variants), and solid points and lines connect the averaged binding values for each genotype. Red-orange lines highlight the impact of introducing the Q493E mutation in different sequence backgrounds. Asterisk indicates expected triple-mutant binding affinity assuming additivity of the single-mutant effects as measured in the BA.2.86 wildtype background. (**C**) Subset of the sarbecovirus RBD sequence alignment showing unique combinations of residues at positions 455, 456, and 493 that have evolved across different sarbecoviruses. Sequence names are colored according to RBD phylogenetic clade as in (Starr, Zepeda, et al. 2022).

We have previously described strong epistatic interactions between position 493 with positions 455 and 456 (Taylor & Starr 2023), and so we wondered whether the deleterious effect of Q493E was modulated by the co-occurring L455S and/or F456L mutations. To address this question, we measured ACE2 binding in isogenic yeast-display titration assays for the L455S, F456L, and Q493E mutations in single, double, and triple mutant combinations (**Figure 4B**). We found that the Q493E mutation is affinity-decreasing on all backgrounds tested except when sampled together with both L455S and F456L, where the effect of Q493E switches in sign to be affinity-enhancing. This result is consistent with recent surface plasmon resonance data (Jian et al. 2024). This strong positive epistasis between L455S, F456L, and Q493E allows KP.3 to maintain ACE2 binding affinity between the levels of JN.1 and BA.2.86, which coupled with its increased immune evasiveness compared to JN.1 (Jian et al. 2024; Kaku, Yo, et al. 2024), likely explains its current rise in frequency.

This second observation of strong epistasis between positions 455, 456, and 493 (Taylor & Starr 2023) is intriguing in light of the large sequence variation at these positions that occurs over broader sarbecovirus evolution (**Figure 4C**). Of note, the GX-Pangolin virus, which can bind strongly and enter efficiently via human ACE2 (Zhang et al. 2021), also naturally samples the Q493E mutation despite sharing the L455 and F456 residues with ancestral SARS-CoV-2 variants like Wuhan-Hu-1 where Q493E is highly detrimental. The presence of Q493E in the GX-Pangolin RBD may therefore point to alternative epistatic interactions that can also compensate for Q493E outside of changes to residues 455 or 456, highlighting the plastic nature of mutational effects in sarbecovirus RBDs.

## DISCUSSION

Here we report deep mutational scanning measurements of the impact of mutations in the BA.2.86 RBD on ACE2 binding and folded RBD expression. We anticipate ongoing utility of these mutational maps in evolutionary forecasting and surveillance of SARS-CoV-2 evolution (Cao et al. 2023; Dadonaite et al. 2024; Greaney, Starr, & Bloom 2022; Maher et al. 2022).

Our measurements reveal only modest epistatic shifts in mutational effects between BA.2 and BA.2.86 despite a similar number of substitutions separating these variants as separates BA.2 from the ancestral Wuhan-Hu-1 RBD. The strong and widespread epistatic shifts caused by N501Y during early SARS-CoV-2 variant evolution therefore continues to be an anomaly in its scale (Starr, Greaney, Hannon, et al. 2022). Nonetheless, small-scale epistatic shifts have also proven important in variant emergence, where unexpected mutations like Q493E (which contributes to KP.3’s enhanced antibody escape (Jian et al. 2024; Kaku, Yo, et al. 2024)) remain capable of occurring on variant backgrounds that epistatically optimize their functional impacts.

We have now described two different sets of sites where the impacts of mutations are highly context dependent. The first, demonstrated here and in prior publication (Jian et al. 2023, 2024; Taylor & Starr 2023), comprises residues 455, 456, and 493 in the central beta-strand of the RBD ACE2-contact surface. The other, demonstrated by us and others in prior work (Moulana et al. 2022; Starr, Greaney, Hannon, et al. 2022; Zahradník et al. 2021), comprises residues 498 and 501. All five of these residues are hotspots of evolution in SARS-CoV-2 as well as the broader sarbecovirus lineage, but the strong epistasis among these positions likely limits our ability to predict what new combinations are compatible with human ACE2 binding. Future work to exhaustively characterize epistatic interaction among the entire suite of amino acids that can occur at these positions would therefore have great utility for ongoing evolutionary modeling of SARS-CoV-2 and sarbecovirus evolution.

## MATERIALS & METHODS

### Mutant libraries

We cloned yeast codon-optimized RBD sequences (amino acids N331 – T531 by Wuhan-Hu-1 reference numbering, the numbering index we use throughout the manuscript) from Omicron BA.2.86 into a yeast surface display plasmid as described (Starr, Greaney, Stewart, et al. 2022). Parental plasmid and associated sequence map is available from Addgene (Addgene ID 222231) and https://github.com/tstarrlab/SARS-CoV-2-RBD_DMS_Omicron-EG5-FLip-BA286/blob/main/data/297_pETcon_SARS2_Omicron-BA286.gb.

Site-saturation mutagenesis libraries spanning all 200 positions in the BA.2.86 RBD were produced by Twist Bioscience. We programmed the introduction of precise codon mutations to encode the 19 possible amino acid mutations at each RBD position and a single-codon deletion. To ensure an adequate level of relevant control variants in the library, stop codon mutations were programmed to be introduced at every other position for the first 50 positions, and wildtype codons were specified at every other position for the first 100 positions. Libraries were delivered as dsDNA oligonucleotides with constant flanking sequences. The “mutant RBD fragment” sequence delivered for BA.2.86 (where uppercase letters denote mutated region) is:

~~~
tctgcaggctagtggtggaggaggctctggtggaggcggccgcggaggcggagggtcggctagccatatgA
ACGTTACCAACTTGTGTCCATTCCATGAAGTTTTCAATGCTACTAGATTCGCTTCTGTTTACGCTTGGAATAGAACT
AGAATCTCTAACTGCGTTGCTGACTATTCTGTCTTGTACAATTTTGCTCCATTCTTCGCTTTCAAGTGCTATGGTGT
TTCTCCAACTAAGTTGAACGATTTGTGTTTCACCAACGTTTACGCCGATTCCTTTGTTATTAAAGGTAACGAAGTCT
CCCAAATTGCTCCAGGTCAAACTGGTAATATTGCCGATTACAATTACAAGTTGCCAGATGATTTCACCGGTTGTGTT
ATTGCTTGGAACTCTAACAAGTTGGATTCTAAGCATTCTGGCAACTACGATTACTGGTACAGGTTGTTCCGTAAGTC
CAAATTGAAGCCATTCGAAAGAGATATTTCCACCGAAATCTATCAAGCTGGTAACAAGCCATGTAAAGGTAAAGGTC
CAAACTGTTACTTCCCATTGCAATCTTACGGTTTCAGACCAACTTATGGTGTTGGTCATCAACCATACAGAGTTGTT
GTTTTGTCTTTCGAGTTGTTGCATGCTCCAGCTACTGTTTGTGGTCCAAAGAAATCTACTctcgaggggggcggttc
cgaacaaaagcttatttctgaagaggacttgtaatagagatctgataacaacagtgtagatgtaacaaaatcgactt
tgttcccactgtacttttagctcg
~~~

A second dsDNA fragment encoding constant flanks and a randomized N16 barcode was produced via PCR off of the parental vector with primer-based sequence additions (primers described in (Greaney, Starr, Eguia, et al. 2022; Starr et al. 2020)). This “barcode fragment” sequence is:

~~~
cgactttgttcccactgtacttttagctcgtacaaaatacaatatacttttcatttctccgtaaacaacat
gttttcccatgtaatatccttttctatttttcgttccgttaccaactttacacatactttatatagctattcacttc
tatacactaaaaaactaagacaattttaattttgctgcctgccatatttcaatttgttataaattcctataatttat
cctattagtagctaaaaaaagatgaatgtgaatcgaatcctaagagaattaatgatacggcgaccaccgagatctac
actctttccctacacgacgctcttccgatctNNNNNNNNNNNNNNNNgcggccgcgagctccaattcgccctatagt
gagtcgtattacaattcactgg
~~~

The mutant RBD fragment and barcode fragment were combined with NotI/SacI-digested parental plasmid backbone via HiFi Assembly. The structure of the final assembled library plasmid is available on GitHub: https://github.com/tstarrlab/SARS-CoV-2-RBD_DMS_Omicron-EG5-FLip-BA286/blob/main/data/297lib_pETcon_SARS2_Omicron-BA286.gb. Assembled library plasmids were electroporated into *E. coli* (NEB 10-beta, New England Biolabs C3020K), and plated at limiting dilutions on LB+ampicillin plates. For each library, duplicate plates corresponding to an estimated bottleneck of ∼80,000 cfu were scraped and plasmid purified, such that each of the 4000 RBD mutations are linked to an average of 20 barcodes. For positions that failed mutagenesis QC from Twist, in-house mutagenesis pools for each position were constructed via PCR with NNS mutagenic primers, Gibson assembled, plated to approximately ∼400 cfu per position, and plasmid purified and pooled with the primary plasmid library. Plasmid libraries are available from Addgene (Addgene ID 1000000248). Plasmid libraries were transformed into the AWY101 yeast strain (Wentz & Shusta 2007) at 10-μg scale according to the protocol of Gietz and Schiestl (Gietz & Schiestl 2007), and aliquots of 9 OD*mL of yeast outgrowth were flash frozen and stored at -80°C.

As described previously (Greaney, Starr, Eguia, et al. 2022; Starr et al. 2020; Starr, Greaney, Hannon, et al. 2022; Starr, Greaney, Stewart, et al. 2022), we sequenced NotI-digested plasmid libraries on a PacBio Sequel IIe to generate long sequence reads spanning the N16 barcode and mutant RBD coding sequence. The resulting circular consensus sequence (CCS) reads are available on the NCBI Sequence Read Archive (SRA), BioProject PRJNA770094, BioSample SAMN42557482. PacBio CCSs were processed using alignparse version 0.6.0 (Crawford & Bloom 2019) to call N16 barcode sequence and RBD variant genotype and filter for high-quality sequences. Analysis of the PacBio sequencing indicates that all but 4 of the intended 4000 RBD mutations were sampled on ≥1 barcode in the BA.2.86 libraries, with even coverage (Figure 1D). Complete computational pipelines and summary plots for PacBio data processing and library analysis are available on GitHub: https://github.com/tstarrlab/SARS-CoV-2-RBD_DMS_Omicron-EG5-FLip-BA286/blob/main/results/summary/process_ccs_BA286.md. Final barcode-variant lookup table is available on GitHub: https://github.com/tstarrlab/SARS-CoV-2-RBD_DMS_Omicron-EG5-FLip-BA286/blob/main/results/variants/codon_variant_table_BA286.csv.

Note, we constructed and assayed site-saturation mutagenesis libraries in the EG.5 and FLip (i.e. HK.3) variant backgrounds in parallel with the BA.2.86 library. However, both of these libraries exhibited certain pathologies in the subsequent deep mutational scanning experiments, notably the presence of hydrophobic mutations at the ACE2 interface (e.g., N417Y, A475L, N487Y, among others) that obtained dubiously high ACE2-binding affinity values. We noticed that these libraries also had higher background staining with our flow cytometry fluorophores, which suggested a degree of nonspecific binding or stickiness, though we did not follow up on these observations. We do still report the data on EG.5 and FLip in the GitHub repository linked throughout these Methods, but only report the high-quality BA.2.86 data in this manuscript and in our public-facing datasets (e.g. interactive websites). EG.5 and FLip data are likely of value but should be interpreted with caution.

### Deep mutational scanning for ACE2-binding affinity

The effects of mutations on ACE2 binding affinity were determined via FACS-seq assays (Adams et al. 2016; Starr et al. 2020; Starr, Greaney, Hannon, et al. 2022; Starr, Greaney, Stewart, et al. 2022; Taylor & Starr 2023). Titrations were performed in duplicate on independently barcoded mutant libraries. Frozen yeast libraries were thawed, grown overnight at 30°C in SD -Ura -Trp media (8 g/L Yeast Nitrogen Base, 2 g/L -Ura -Trp Synthetic Complete dropout powder and 2% w/v dextrose), and backdiluted to 0.67 OD600 in SG -Ura -Trp + 0.1%D (recipe as above but with 2% galactose and 0.1% dextrose in place of the 2% dextrose) to induce RBD expression, which proceeded for 22-24 hours at 19°C with gentle mixing.

Induced cells were washed with PBS-BSA (BSA 0.2 mg/L), split into 16-OD*mL aliquots, and incubated with biotinylated monomeric human ACE2 protein (ACROBiosystems AC2-H82E8) across a concentration range from 10^−6^ to 10^−13^ M at 1-log intervals, plus a 0 M sample. Incubations equilibrated overnight at room temperature with gentle mixing. Yeast were washed twice with ice-cold PBS-BSA and fluorescently labeled for 1 hr at 4°C with 1:100 FITC-conjugated chicken anti-Myc (Immunology Consultants CMYC-45F) to detect yeast-displayed RBD protein and 1:200 PE-conjugated streptavidin (Thermo Fisher S866) to detect bound ACE2. Cells were washed and resuspended in PBS-BSA for flow cytometry.

At each ACE2 sample concentration, single RBD^+^ cells were partitioned into bins of ACE2 binding (PE fluorescence) as shown in Figure 1E using a BD FACSAria II. A minimum of 10 million cells were collected at each sample concentration, and sorted into SD -Ura -Trp with pen-strep antibiotic and 1% BSA. Collected cells in each bin were grown overnight in 1 mL SD -Ura -Trp + pen-strep, and plasmid was isolated using a 96-well yeast miniprep kit (Zymo D2005) according to kit instructions, with the addition of an extended (>2 hr) Zymolyase treatment and a -80°C freeze/thaw prior to cell lysis. N16 barcodes in each post-sort sample were PCR amplified as described in (Starr et al. 2020) and submitted for Illumina sequencing. Barcode sequence reads are available on the NCBI SRA, BioProject PRJNA770094, BioSample SAMN42557522.

Demultiplexed Illumina barcode reads were matched to library barcodes in barcode-mutant lookup tables using dms_variants (version 1.4.3), yielding a table of counts of each barcode in each FACS bin, available at https://github.com/tstarrlab/SARS-CoV-2-RBD_DMS_Omicron-EG5-FLip-BA286/blob/main/results/counts/variant_counts.csv. Read counts in each FACS bin were downweighted by the ratio of total sequence reads from a bin to the number of cells that were sorted into that bin from the FACS log.

We estimated the level of ACE2 binding of each barcoded mutant at each ACE2 concentration based on its distribution of counts across FACS bins as the simple mean bin (Starr et al. 2020). We determined the ACE2-binding constant *K*_D_ for each barcoded mutant via nonlinear least-squares regression using the standard non-cooperative Hill equation relating the mean sort bin to the ACE2 labeling concentration and free parameters *a* (titration response range) and *b* (titration curve baseline):

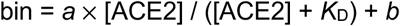

The measured mean bin value for a barcode at a given ACE2 concentration was excluded from curve fitting if fewer than 2 counts were observed across the four FACS bins or if counts exhibited bimodality (>40% of counts of a barcode were found in each of two non-consecutive bins). To avoid errant fits, we constrained the value *b* to (1, 1.5), *a* to (2, 3), and *K*_D_ to (10^−15^, 10^−5^). The fit for a barcoded variant was discarded if the average cell count across all sample concentrations was below 2, or if more than one sample concentration was missing. We also discarded curve fits where the normalized mean square residual (residuals normalized relative to the fit response parameter *a*) was >20 times the median value across all titration fits. Final binding constants were expressed as -log_10_(*K*_D_), where higher values indicate higher binding affinity. The complete computational pipeline for calculating and filtering per-barcode binding constants is available at https://github.com/tstarrlab/SARS-CoV-2-RBD_DMS_Omicron-EG5-FLip-BA286/blob/main/results/summary/compute_binding_Kd.md, and per-barcode affinity values are available at https://github.com/tstarrlab/SARS-CoV-2-RBD_DMS_Omicron-EG5-FLip-BA286/blob/main/results/binding_Kd/bc_binding.csv.

The affinity measurements of replicate barcodes representing an identical amino acid mutant were averaged within each experimental duplicate. The correlations in collapsed affinities in each duplicate experiment are shown in Figure 1G. The final measurement was determined as the average of duplicate measurements. The final -log_10_(*K*_D_) for each mutant and number of replicate barcode collapsed into this final measurement for each RBD mutant are given in Supplemental Data 1 and https://github.com/tstarrlab/SARS-CoV-2-RBD_DMS_Omicron-EG5-FLip-BA286/blob/main/results/final_variant_scores/final_variant_scores.csv, which includes data also from prior SARS-CoV-2 variant DMS datasets (Starr, Greaney, Hannon, et al. 2022; Starr, Greaney, Stewart, et al. 2022; Taylor & Starr 2023).

### RBD expression deep mutational scanning

Pooled libraries were grown and induced for RBD expression as described above. Induced cells were washed and labeled with 1:100 FITC-conjugated chicken anti-Myc to label for RBD expression via a C-terminal Myc tag, and washed in preparation for FACS. Single cells were partitioned into bins of RBD expression (FITC fluorescence) using a BD FACSAria II as shown in Figure 1F. A total of >18 million viable cells (estimated by plating dilutions of post-sort samples) were collected across bins for each library. Cells in each bin were grown out in SD -Ura -Trp + pen-strep + 1% BSA, plasmid isolated, and N16 barcodes sequenced as described above. Barcode reads are available on the NCBI SRA, BioProject PRJNA770094, BioSample SAMN42557522.

Demultiplexed Illumina barcode reads were matched to library barcodes in barcode-mutant lookup tables using dms_variants (version 0.8.9), yielding a table of counts of each barcode in each FACS bin, available at https://github.com/tstarrlab/SARS-CoV-2-RBD_DMS_Omicron-EG5-FLip-BA286/blob/main/results/counts/variant_counts.csv. Read counts in each bin were downweighted using the post-sort colony counts instead of the FACS log counts as with ACE2 titrations above to account for unequal viability of cells in FITC fluorescence bins (i.e., many cells in bin 1 are non-expressing because they have lost the low-copy expression plasmid and do not grow out post-FACS in selective media).

We estimated the level of RBD expression (log-mean fluorescence intensity, logMFI) of each barcoded mutant based on its distribution of counts across FACS bins and the known log-transformed fluorescence boundaries of each sort bin using a maximum likelihood approach (Peterman & Levine 2016; Starr et al. 2020) implemented via the fitdistrplus package in R (Delignette-Muller & Dutang 2015). Expression measurements were discarded for barcodes for which fewer than 10 counts were observed across the four FACS bins. The full pipeline for computing per-barcode expression values is available at https://github.com/tstarrlab/SARS-CoV-2-RBD_DMS_Omicron-EG5-FLip-BA286/blob/main/results/summary/compute_expression_meanF.md, and per-barcode expression measurements are available at https://github.com/tstarrlab/SARS-CoV-2-RBD_DMS_Omicron-EG5-FLip-BA286/blob/main/results/expression_meanF/bc_expression.csv. Final mutant expression values were collapsed within and across replicates as described above, with correlation between experimental replicates shown in Figure 1H. Final mutant expression values and number of replicate barcode collapsed into this final measurement for each RBD mutant are available in Supplemental Data 1 and at https://github.com/tstarrlab/SARS-CoV-2-RBD_DMS_Omicron-EG5-FLip-BA286/blob/main/results/final_variant_scores/final_variant_scores.csv.

### Quantification of epistasis

Epistatic shifts at each site between pairs of RBD variants were quantified as described by (Starr, Greaney, Hannon, et al. 2022). Briefly, affinity phenotypes of each mutant at a site were transformed to a probability analog via a Boltzmann weighting, and the “epistatic shift” metric was calculated as the Jensen-Shannon divergence between the vectors of 21 amino acid probabilities (including the deletion character, when present). The Jensen-Shannon divergence ranges from 0 for two vectors of probabilities that are identical to 1 for two vectors that are completely dissimilar. To avoid noisier measurements artifactually inflating the epistatic shift metric, a given amino acid mutation was only included in the computation if it was sampled with a minimum of 3 replicate barcodes in each RBD background being compared. The calculation of epistatic shifts can be found at https://github.com/tstarrlab/SARS-CoV-2-RBD_DMS_Omicron-EG5-FLip-BA286/blob/main/results/summary/epistatic_shifts.md.

### Sites of strong antibody escape

Sites of strong antibody escape are determined from a large aggregation of deep mutational scanning antibody-escape datasets summarized in (Greaney, Starr, & Bloom 2022), available from https://github.com/jbloomlab/SARS2_RBD_Ab_escape_maps (note that this repository has since been replaced by a newer repository integrating additional antibody-escape deep mutational scanning datasets gathered in more updated Omicron backgrounds). We downloaded the aggregate dataset on February 16, 2024, and highlighted sites as “sites of strong escape” if their normalized site-wise escape score averaged across all mAbs was greater than 0.125.

### Isogenic validation of epistasis

Epistasis was validated via ACE2 binding assays with isogenic yeast expressing clonal RBD variants. Single, double, and triple mutants in the BA.2.86 background were constructed by Twist Bioscience and individually transformed into the AWY101 yeast strain. RBD expression was induced as above, and induced cells (0.083 OD*mL per sample) were incubated across a concentration series from 10^−6^ to 10^−12^M (plus a 0M sample) monomeric human ACE2 in 100μL in 96-well V-bottom plates, washed, and labeled for RBD expression and ACE2 binding. ACE2 binding (log-geometric mean of PE fluorescence) was determined among RBD^+^ cells via flow cytometry using a BD FACSCelesta. ACE2 binding versus concentration was fit to a sigmoidal curve in R using nonlinear least squares regression, with the midpoint reported as the EC_50_. Each measurement was made in duplicate.

## Acknowledgements

We thank Jesse Bloom and Bernadeta Dadonaite for helpful discussion. We thank the Flow Cytometry Core Facility at the University of Utah Health Sciences Campus supported by NIH (5P30CA042014-24), and the University of Utah Center for High Performance Computing supported by NIH (1S10OD021644-01A1).

## Data and Materials Availability

Site saturation mutagenesis libraries and respective isogenic parental plasmid stocks are available from Addgene (Addgene ID 1000000248). Raw sequencing data are on the NCBI SRA under BioProject PRJNA770094, BioSamples SAMN42557482 (PacBio sequencing) and SAMN42557522 (Illumina barcode sequencing for ACE2 binding and expression experiments). All code and data at various stages of processing is available at https://github.com/tstarrlab/SARS-CoV-2-RBD_DMS_Omicron-EG5-FLip-BA286/tree/main. Outlines of the analytical pipelines and links to descriptive notebooks for each analytical step are available at https://github.com/tstarrlab/SARS-CoV-2-RBD_DMS_Omicron-EG5-FLip-BA286/blob/main/results/summary/summary.md. Final mutant deep mutational scanning measurements are available in Supplemental Data 1, and interactive visualizations of key data are available at: https://tstarrlab.github.io/SARS-CoV-2-RBD_DMS_Omicron-EG5-FLip-BA286/.

## REFERENCES

Adams, R. M., Mora, T., Walczak, A. M., & Kinney, J. B. (2016). ‘Measuring the sequence-affinity landscape of antibodies with massively parallel titration curves’, eLife, 5. DOI: 10.7554/eLife.23156

Cao, Y., Jian, F., Wang, J., Yu, Y., Song, W., Yisimayi, A., Wang, J., et al. (2023). ‘Imprinted SARS-CoV-2 humoral immunity induces convergent Omicron RBD evolution’, Nature, 614/7948: 521–9.

Cao, Y., Wang, J., Jian, F., Xiao, T., Song, W., Yisimayi, A., Huang, W., et al. (2022). ‘Omicron escapes the majority of existing SARS-CoV-2 neutralizing antibodies’, Nature, 602/7898: 657–63. Nature Publishing Group.

Carabelli, A. M., Peacock, T. P., Thorne, L. G., Harvey, W. T., Hughes, J., COVID-19 Genomics UK Consortium, Peacock, S. J., et al. (2023). ‘SARS-CoV-2 variant biology: immune escape, transmission and fitness’, Nature reviews. Microbiology. DOI: 10.1038/s41579-022-00841-7

Crawford, K. H. D., & Bloom, J. D. (2019). ‘alignparse: A Python package for parsing complex features from high-throughput long-read sequencing’, Journal of open source software, 4/44. DOI: 10.21105/joss.01915

Dadonaite, B., Brown, J., McMahon, T. E., Farrell, A. G., Figgins, M. D., Asarnow, D., Stewart, C., et al. (2024). ‘Spike deep mutational scanning helps predict success of SARS-CoV-2 clades’, Nature, 631/8021: 617–26. Springer Science and Business Media LLC.

Delignette-Muller, M., & Dutang, C. (2015). ‘fitdistrplus: An R Package for Fitting Distributions’, Journal of Statistical Software, Articles, 64/4: 1–34.

Ellis, D., Brunette, N., Crawford, K. H. D., Walls, A. C., Pham, M. N., Chen, C., Herpoldt, K.-L., et al. (2021). ‘Stabilization of the SARS-CoV-2 Spike Receptor-Binding Domain Using Deep Mutational Scanning and Structure-Based Design’, Frontiers in immunology, 12: 710263.

Fowler, D. M., & Fields, S. (2014). ‘Deep mutational scanning: a new style of protein science’, Nature methods, 11/8: 801–7.

Francino-Urdaniz, I. M., Steiner, P. J., Kirby, M. B., Zhao, F., Haas, C. M., Barman, S., Rhodes, E. R., et al. (2021). ‘One-shot identification of SARS-CoV-2 S RBD escape mutants using yeast screening’, Cell reports, 36/9: 109627.

Gietz, R. D., & Schiestl, R. H. (2007). ‘Large-scale high-efficiency yeast transformation using the LiAc/SS carrier DNA/PEG method’, Nature protocols, 2/1: 38–41.

Greaney, A. J., Starr, T. N., & Bloom, J. D. (2022). ‘An antibody-escape estimator for mutations to the SARS-CoV-2 receptor-binding domain’, Virus evolution, 8/1: veac021.

Greaney, A. J., Starr, T. N., Eguia, R. T., Loes, A. N., Khan, K., Karim, F., Cele, S., et al. (2022). ‘A SARS-CoV-2 variant elicits an antibody response with a shifted immunodominance hierarchy’, PLoS pathogens, 18/2: e1010248. Public Library of Science.

Greaney, A. J., Starr, T. N., Gilchuk, P., Zost, S. J., Binshtein, E., Loes, A. N., Hilton, S. K., et al. (2021). ‘Complete Mapping of Mutations to the SARS-CoV-2 Spike Receptor-Binding Domain that Escape Antibody Recognition’, Cell host & microbe, 29/1: 44–57.e9.

Jian, F., Feng, L., Yang, S., Yu, Y., Wang, L., Song, W., Yisimayi, A., et al. (2023). ‘Convergent evolution of SARS-CoV-2 XBB lineages on receptor-binding domain 455-456 synergistically enhances antibody evasion and ACE2 binding’, PLoS pathogens, 19/12: e1011868.

Jian, F., Wang, J., Yisimayi, A., Song, W., Xu, Y., Chen, X., Niu, X., et al. (2024). Evolving antibody response to SARS-CoV-2 antigenic shift from XBB to JN.1. bioRxiv. DOI: 10.1101/2024.04.19.590276

Kaku, Y., Uriu, K., Okumura, K., The Genotype to Phenotype Japan (G2P-Japan) Consortium, Ito, J., & Sato, K. (2024). Virological characteristics of the SARS-CoV-2 KP.3.1.1 variant. bioRxiv. DOI: 10.1101/2024.07.16.603835

Kaku, Y., Yo, M. S., Tolentino, J. E., Uriu, K., Okumura, K., Genotype to Phenotype Japan (G2P-Japan) Consortium, Ito, J., et al. (2024). ‘Virological characteristics of the SARS-CoV-2 KP.3, LB.1, and KP.2.3 variants’, The Lancet infectious diseases. DOI: 10.1016/S1473-3099(24)00415-8

Khan, K., Lustig, G., Römer, C., Reedoy, K., Jule, Z., Karim, F., Ganga, Y., et al. (2023). ‘Evolution and neutralization escape of the SARS-CoV-2 BA.2.86 subvariant’, Nature communications, 14/1: 8078.

Kowalski, J. M., Parekh, R. N., & Wittrup, K. D. (1998). ‘Secretion efficiency in Saccharomyces cerevisiae of bovine pancreatic trypsin inhibitor mutants lacking disulfide bonds is correlated with thermodynamic stability’, Biochemistry, 37/5: 1264–73.

Kugathasan, R., Sukhova, K., Moshe, M., Kellam, P., & Barclay, W. (2023). ‘Deep mutagenesis scanning using whole trimeric SARS-CoV-2 spike highlights the importance of NTD-RBD interactions in determining spike phenotype’, PLoS pathogens, 19/8: e1011545.

Liu, C., Zhou, D., Dijokaite-Guraliuc, A., Supasa, P., Duyvesteyn, H. M. E., Ginn, H. M., Selvaraj, M., et al. (2024). ‘A structure-function analysis shows SARS-CoV-2 BA.2.86 balances antibody escape and ACE2 affinity’, Cell reports. Medicine, 5/5: 101553.

Maher, M. C., Bartha, I., Weaver, S., di Iulio, J., Ferri, E., Soriaga, L., Lempp, F. A., et al. (2022). ‘Predicting the mutational drivers of future SARS-CoV-2 variants of concern’, Science translational medicine, eabk3445.

Ma, W., Fu, H., Jian, F., Cao, Y., & Li, M. (2023). ‘Immune evasion and ACE2 binding affinity contribute to SARS-CoV-2 evolution’, Nature ecology & evolution, 7/9: 1457–66.

Moulana, A., Dupic, T., Phillips, A. M., Chang, J., Nieves, S., Roffler, A. A., Greaney, A. J., et al. (2022). ‘Compensatory epistasis maintains ACE2 affinity in SARS-CoV-2 Omicron BA.1’, Nature communications, 13/1: 7011.

Ouyang, W. O., Tan, T. J. C., Lei, R., Song, G., Kieffer, C., Andrabi, R., Matreyek, K. A., et al. (2022). ‘Probing the biophysical constraints of SARS-CoV-2 spike N-terminal domain using deep mutational scanning’, Science advances, 8/47: eadd7221.

Peterman, N., & Levine, E. (2016). ‘Sort-seq under the hood: implications of design choices on large-scale characterization of sequence-function relations’, BMC genomics, 17: 206.

Shusta, E. V., Kieke, M. C., Parke, E., Kranz, D. M., & Wittrup, K. D. (1999). ‘Yeast polypeptide fusion surface display levels predict thermal stability and soluble secretion efficiency’, Journal of molecular biology, 292/5: 949–56.

Starr, T. N., Greaney, A. J., Addetia, A., Hannon, W. W., Choudhary, M. C., Dingens, A. S., Li, J. Z., et al. (2021). ‘Prospective mapping of viral mutations that escape antibodies used to treat COVID-19’, Science, 371/6531: 850–4. American Association for the Advancement of Science.

Starr, T. N., Greaney, A. J., Dingens, A. S., & Bloom, J. D. (2021). ‘Complete map of SARS-CoV-2 RBD mutations that escape the monoclonal antibody LY-CoV555 and its cocktail with LY-CoV016’, Cell reports. Medicine, 2/4: 100255.

Starr, T. N., Greaney, A. J., Hannon, W. W., Loes, A. N., Hauser, K., Dillen, J. R., Ferri, E., et al. (2022). ‘Shifting mutational constraints in the SARS-CoV-2 receptor-binding domain during viral evolution’, Science, 377/6604: 420–4.

Starr, T. N., Greaney, A. J., Hilton, S. K., Ellis, D., Crawford, K. H. D., Dingens, A. S., Navarro, M. J., et al. (2020). ‘Deep Mutational Scanning of SARS-CoV-2 Receptor Binding Domain Reveals Constraints on Folding and ACE2 Binding’, Cell, 182/5: 1295–310.e20.

Starr, T. N., Greaney, A. J., Stewart, C. M., Walls, A. C., Hannon, W. W., Veesler, D., & Bloom, J. D. (2022). ‘Deep mutational scans for ACE2 binding, RBD expression, and antibody escape in the SARS-CoV-2 Omicron BA.1 and BA.2 receptor-binding domains’, PLoS pathogens, 18/11: e1010951.

Starr, T. N., & Thornton, J. W. (2016). ‘Epistasis in protein evolution’, Protein science: a publication of the Protein Society, 25/7: 1204–18.

Starr, T. N., Zepeda, S. K., Walls, A. C., Greaney, A. J., Alkhovsky, S., Veesler, D., & Bloom, J. D. (2022). ‘ACE2 binding is an ancestral and evolvable trait of sarbecoviruses’, Nature, 603/7903: 913–8. Nature Publishing Group.

Taft, J. M., Weber, C. R., Gao, B., Ehling, R. A., Han, J., Frei, L., Metcalfe, S. W., et al. (2022). ‘Deep mutational learning predicts ACE2 binding and antibody escape to combinatorial mutations in the SARS-CoV-2 receptor-binding domain’, Cell, 185/21: 4008–22.e14.

Tamura, T., Mizuma, K., Nasser, H., Deguchi, S., Padilla-Blanco, M., Oda, Y., Uriu, K., et al. (2024). ‘Virological characteristics of the SARS-CoV-2 BA.2.86 variant’, Cell host & microbe, 32/2: 170–80.e12.

Taylor, A. L., & Starr, T. N. (2023). ‘Deep mutational scans of XBB.1.5 and BQ.1.1 reveal ongoing epistatic drift during SARS-CoV-2 evolution’, PLoS pathogens, 19/12: e1011901.

Toelzer, C., Gupta, K., Yadav, S. K. N., Borucu, U., Davidson, A. D., Kavanagh Williamson, M., Shoemark, D. K., et al. (2020). ‘Free fatty acid binding pocket in the locked structure of SARS-CoV-2 spike protein’, Science, 370/6517: 725–30.

Wang, Q., Guo, Y., Liu, L., Schwanz, L. T., Li, Z., Nair, M. S., Ho, J., et al. (2023). ‘Antigenicity and receptor affinity of SARS-CoV-2 BA.2.86 spike’, Nature, 624/7992: 639–44.

Wentz, A. E., & Shusta, E. V. (2007). ‘A novel high-throughput screen reveals yeast genes that increase secretion of heterologous proteins’, Applied and environmental microbiology, 73/4: 1189–98.

Yang, S., Yu, Y., Jian, F., Song, W., Yisimayi, A., Chen, X., Xu, Y., et al. (2023). ‘Antigenicity and infectivity characterisation of SARS-CoV-2 BA.2.86’, The Lancet infectious diseases, 23/11: e457–9.

Yang, S., Yu, Y., Xu, Y., Jian, F., Song, W., Yisimayi, A., Wang, P., et al. (2024). ‘Fast evolution of SARS-CoV-2 BA.2.86 to JN.1 under heavy immune pressure’, The Lancet infectious diseases, 24/2: e70–2.

Zahradník, J., Marciano, S., Shemesh, M., Zoler, E., Harari, D., Chiaravalli, J., Meyer, B., et al. (2021). ‘SARS-CoV-2 variant prediction and antiviral drug design are enabled by RBD in vitro evolution’, Nature Microbiology, 6/9: 1188–98. Nature Publishing Group.

Zhang, S., Qiao, S., Yu, J., Zeng, J., Shan, S., Tian, L., Lan, J., et al. (2021). ‘Bat and pangolin coronavirus spike glycoprotein structures provide insights into SARS-CoV-2 evolution’, Nature communications, 12/1: 1607.

